# Evidence for spreading seizure as a cause of theta-alpha activity electrographic pattern in stereo-EEG seizure recordings

**DOI:** 10.1101/2020.07.22.215442

**Authors:** Viktor Sip, Julia Scholly, Maxime Guye, Fabrice Bartolomei, Viktor Jirsa

## Abstract

Intracranial electroencephalography is a standard tool in clinical evaluation of patients with focal epilepsy. Various early electrographic seizure patterns differing in frequency, amplitude, and waveform of the oscillations are observed. The pattern most common in the areas of seizure propagation is the so-called theta-alpha activity (TAA), whose defining features are oscillations in the *θ* – *α* range and gradually increasing amplitude. A deeper understanding of the mechanism underlying the generation of the TAA pattern is however lacking. In this work we evaluate the hypothesis that the TAA patterns are caused by seizures spreading across the cortex. To do so, we perform simulations of seizure dynamics on detailed patient-derived cortical surfaces using the spreading seizure model as well as reference models with one or two homogeneous sources. We then detect the occurrences of the TAA patterns both in the simulated stereo-electroencephalographic signals and in the signals of recorded epileptic seizures from a cohort of fifty patients, and we compare the features of the groups of detected TAA patterns to assess the plausibility of the different models. Our results show that spreading seizure hypothesis is qualitatively consistent with the evidence available in the seizure recordings, and it can explain the features of the detected TAA groups best among the examined models.

## 1. Introduction

### Intracranial EEG and early electrographic patterns

Intracranial electroencephalography (iEEG) is an essential tool in clinical evaluation of patients with focal drug-resistant epilepsy and its use in neuroscientific research is steadily growing (Bartolomei et al., 2017; Parvizi and Kastner, 2018). The objective of exploration using iEEG is to understand the spatiotemporal organization of the patient’s epilepsy with the goal to perform resective surgery and render the patient seizure free. In epilepsy both electrocorticography (ECoG) using the subdural grids and stereoelectroencephalography (SEEG) using the depth electrodes are widely employed.

The acquired intracranial recordings show variety of early electrographic seizure patterns, i.e. patterns occurring shortly after the appearance of electrographic seizure activity at the observed site. The patterns differ in frequency, amplitude, and waveform of the oscillations and in their temporal evolution. Several studies attempted to classify these patterns in an effort to distinguish between local onset and propagated seizures and to link the early patterns with different pathologies or with the outcome of a resective surgery (Alarcon et al., 1995; Schiller et al., 1998; Doležalová et al., 2013; Perucca et al., 2014; Singh et al., 2015; Jiménez-Jiménez et al., 2015, 2016; Lagarde et al., 2016).

### TAA electrographic pattern

One of the often described patterns is what we call *theta-alpha activity* (TAA): an early ictal pattern characterized by sustained oscillations in the *θ* – *α* range with gradually increasing amplitude (Fig 1). Such pattern was reported under the names “rhythmic ictal transformation” (Alarcon et al., 1995), “sharp activity at ≤ 13Hz” (Perucca et al., 2014), and “theta/alpha sharp activity” (Lagarde et al., 2016). Furthermore, several other studies include similar category of rhythmic theta-alpha activity, although without explicitly mentioning the gradually increasing amplitude (Schiller et al., 1998; Wennberg et al., 2002; Singh et al., 2015). In the seizure onset zone, the TAA pattern was reported as less common compared low-voltage fast activity and low-frequency high-amplitude spikes patterns (Alarcon et al., 1995; Perucca et al., 2014; Lagarde et al., 2016). However, the pattern was commonly associated with the regions of seizure spread (Schiller et al., 1998; Perucca et al., 2014; Singh et al., 2015).

**Figure 1:**
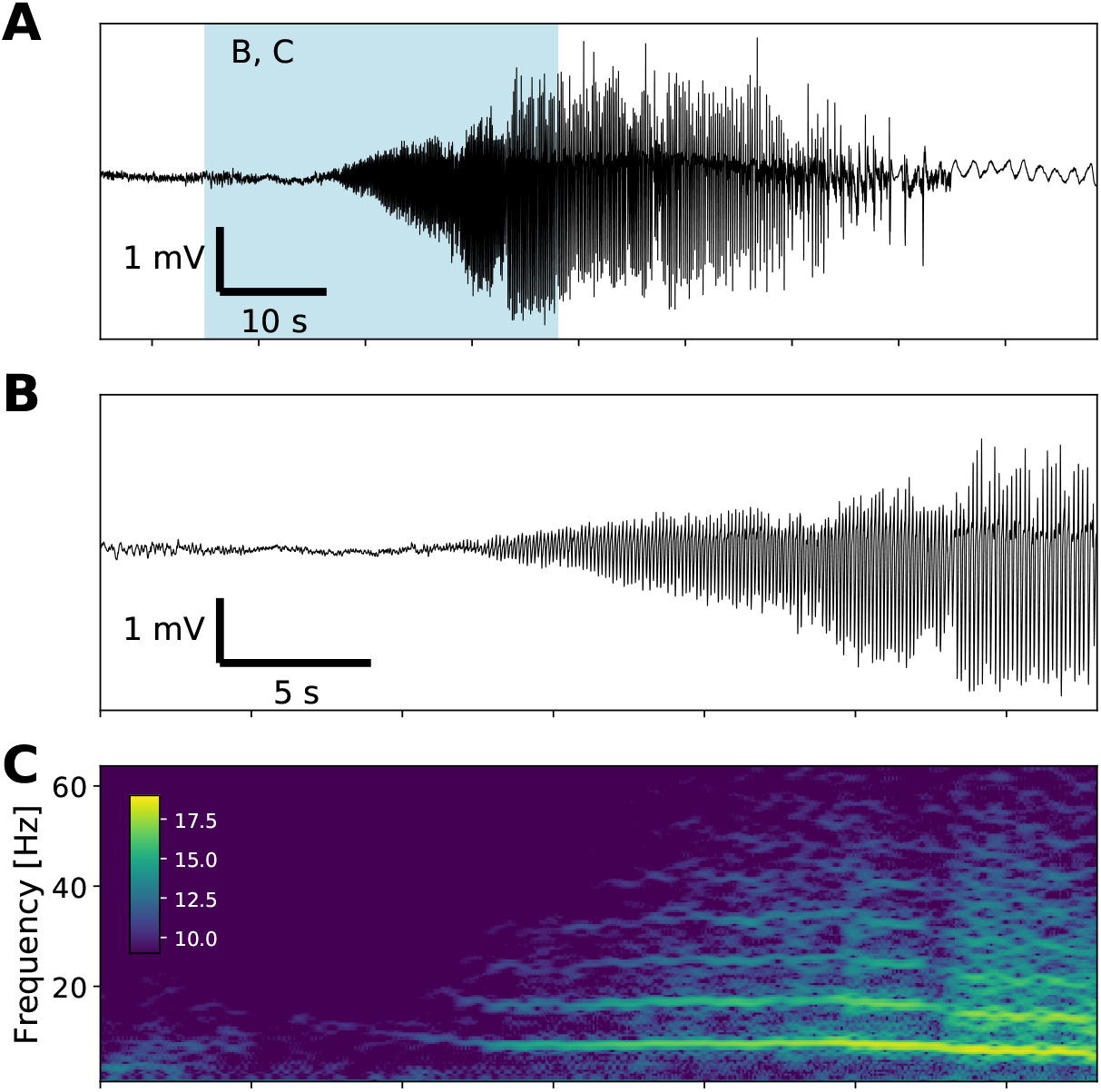
Example of the TAA pattern. (A) Recording of a seizure from a contact of an implanted depth electrode in monopolar representation. Blue background marks the limits of the details in panels B and C. (B) Detail of the TAA pattern at the seizure onset. (C) Time-frequency representation of the signal in panel B (log transformed). The signal is dominated by oscillations around 8 Hz and its higher harmonics.

### Seizures spreading across the cortex

In this work we aim to investigate whether the TAA pattern can be produced by seizures locally spreading across the cortex. This local seizure spread, or propagating ictal wavefront, refers to the scenario where the seizure is initiated at a spatially restricted location (either autonomously or via external intervention), from where it gradually spreads to the surrounding tissue. Such seizure spread was observed in *in vitro* (Wong and Prince, 1990; Trevelyan et al., 2006, 2007) and *in vivo* animal models of epilepsy (Wenzel et al., 2017), as well as in human patients (Schevon et al., 2012; Martinet et al., 2015). The reported velocity of the seizure spread was between 0.1 and 10mm s^−1^, with the exception of situations where the lateral inhibition was suppressed at the time of seizure initiation. In that case the seizure spread velocity was reported to reach up to tens to hundred of mm s^−1^ (Trevelyan et al., 2006, 2007); in this work, we do not model this special case. Theoretical models of epileptic seizure dynamics which reproduce locally spreading seizures exist, and include a network of coupled Wilson-Cowan units (Wang et al., 2017), spatially continuous version of the phenomenological Epileptor model (Proix et al., 2018), or a biophysically-constrained model of seizure dynamics (Liou et al., 2019). The relevance of local propagation for large-scale spatiotemporal organization of seizures in human patients compared to the propagation through the long-range projections (Proix et al., 2017) was not yet determined.

Experimental results indicate that, at least under some conditions, the seizure activity in the recruited tissue has a distinct spatio-temporal organization formed by fast traveling waves of increased firing. Such waves were observed in the cortex of human patients during seizures, spatially extending not only across microelectrode arrays but also across ECoG array, with velocities ranging from 100 to 1000 mm s^−1^ (Wagner et al., 2015; Smith et al., 2016; Martinet et al., 2017). Also this feature can be reproduced by some models of seizure dynamics (Proix et al., 2018; Liou et al., 2019).

### Spreading seizures as a cause of TAA patterns

The two features of spreading seizures - slow wavefront and fast internal traveling waves - motivate our speculations that the locally spreading seizures may be linked to the observed TAA pattern, thus explaining its reported occurrence in the regions of seizure spread. To better elucidate this possible link, one has to first consider the relation of the (unobserved) activity of the neuronal assemblies (to which we will hereafter refer as to the *source* activity) and of the signal recorded by the intracranial electrodes (hereafter referred as the *sensor* activity). The electrodes record the local field potential (LFP) generated by the neuronal assemblies both local and distant. The amplitude of LFP recorded on the sensors is affected both by the synchronization of the neuronal assemblies as well as by the geometry of the cortical tissue and subcortical structures and the exact positions of the contacts of the implanted electrodes (Buzsáki et al., 2012; Herreras, 2016). The LFP performs a spatial averaging of the source activity, and therefore the gradual recruitment of the neuronal tissue by the slowly propagating seizure may manifest itself as a gradually growing amplitude of the oscillations in the recorded signals - the characteristic feature of the TAA pattern. At the same time, if the internal organization of the seizure is dominated by the fast traveling waves, then the firing of the recruited tissue is synchronized by these waves. That, in turn, would give rise to LFP oscillations dominated by the single frequency, which is the second feature of the TAA pattern.

### Goal and organization of the paper

In this work we propose local seizure propagation as a (non-exclusive) candidate mechanism for the generation of electrographic TAA patterns. Here we aim to determine whether this hypothesis is plausible when confronted with the evidence available in the SEEG recordings of epileptic seizures in human patients. To do so, we take the following steps: First, we generate a synthetic data set of SEEG signals by simulating the seizure dynamics on realistic cortical surfaces using model parameters randomly sampled from prescribed parameter range. We use a simple model of spreading seizure as well as reference models of one and two homogeneous sources. Next, using a strict definition of the TAA pattern, we detect these in SEEG recordings obtained from a cohort of fifty subjects, as well as from the simulated SEEG. We then compare the statistical distributions of the features of the TAA patterns detected in the recordings with those detected in the simulated data set in order to assess the plausibility of the propagating seizure model relative to the reference models.

## 2. Results

### Models of seizure activity

For our analysis, we used three models of seizure activity - one homogeneous source, two homogeneous sources, and spreading seizure (Fig. 2). All models follow the same structure. They posit that only a small patch (or two small patches in case of the two sources model) of the cortex is recruited in the seizure activity, and the rest of the cortex is modeled just as a noise source. The source activity is represented on the vertices of the triangulated surface, and differs for the three models: for one homogeneous source, all vertices in the seizure patch follow the same dynamics of oscillations with gradually increasing amplitude (Fig. 2A). In the two homogeneous sources models, there are two patches which are recruited at different times, but the activity in each patch is again homogeneous (Fig. 2B). The patches oscillates with the same frequency, although possibly with different phases. In the spreading seizure model (Fig. 2C), the seizure patch is recruited gradually as the seizure spreads. On the source level, the seizure starts instantly with no transition period. Inside the recruited part of the patch, the activity is organized by fast traveling waves. The simulations are done both with noise-free and noisy seizure activity. For the latter, on top of the deterministic source activity spatially correlated pink noise is added.

**Figure 2:**
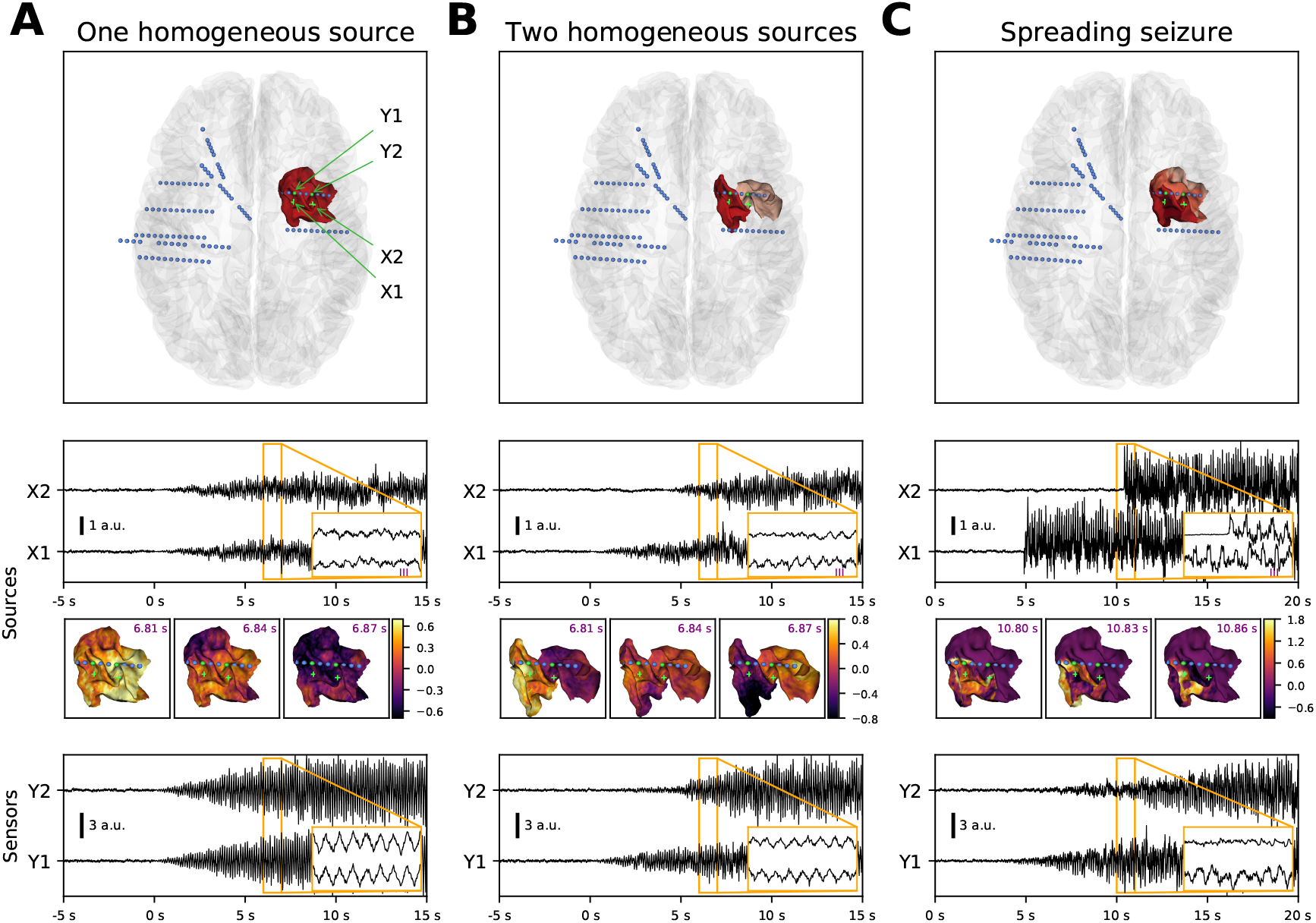
Models of the seizure activity. Each column presents an example of a simulated seizure with the different models in the noisy variation. From top to bottom, the panels show the position of the seizure patch and the implanted electrodes, source activity in two points on the patch, snapshots of the source activity (at time points marked in the inset above), and simulated sensor activity on two contacts located close to the seizure patch. (A) One homogeneous source model posits that the seizure activity is generated by one contiguous cortical patch, where any point follows the same dynamics (apart from the stochastic noise) with gradually increasing oscillations. (B) In the two homogeneous sources model, there are two patches recruited with a delay. On each patch any point follows the same dynamics with gradual onset as in the one source model. (C) In the spreading seizure model the patch is recruited gradually as the seizure spreads (recruitment time is represented by the patch color). At every point, the seizure activity starts instantly with no transition period; the gradual onset on the sensors is caused by the slow spread of the seizure. The lowermost panel shows that all models produce SEEG signal resembling the TAA pattern.

The simulations were performed on the triangulated cortical surfaces obtained from the fifty subjects in the patient cohort. The triangulated surfaces had mean total area 1790.7 cm^2^ (standard deviation 204.6 cm^2^), mean number of vertices 251.9 thousands (s.d. 31.2 thousands), mean number of triangles 503.1 thousands (s.d. 62.4 thousands), mean triangle area 0.356 mm^2^ (s.d. 0.199 mm^2^), and mean edge length 0.924 mm (s.d. 0.295 mm^2^). To assess the influence of the triangulation density, we performed a set of simulations on this standard triangulation and refined triangulation, obtained by splitting every existing triangle into four (Fig. A.7). The results show that using the standard triangulation does not introduce differences of higher order of magnitude than those due to the stochastic background noise, and we thus used the standard triangulation in the rest of the work.

For each subject and each model, 300 simulations were performed, leading to a total of 15000 simulated seizures for each model. For each simulation, the model parameters were drawn randomly from the prescribed parameter range (Tab. 2). The source activity was projected on the sensors using the dipole model of generated local field position. The position of the contacts was derived from patient data and thus constituted a realistic placement of the electrodes. In case that the seizure activity was not detected on any contact in the simulation, new set of parameters was drawn and the simulation was repeated.

### Detection of TAA patterns

Using a cohort of 50 patients with focal epilepsy who underwent clinical evaluation via SEEG, we detected the occurrences of the TAA pattern in the seizure recordings as well as in the simulated SEEG signals (Fig. 3A). These patterns were detected on all channels separately. Next, we restricted our analysis to the TAA patterns occurring on at least four consecutive contacts on a single electrode, which we in the following text call *TAA groups*. This step was motivated by the reasoning that the source activity often affects multiple electrode contacts at once, and that more information about the spatial configuration of the sources can be extracted from the properties of these groups.

**Figure 3:**
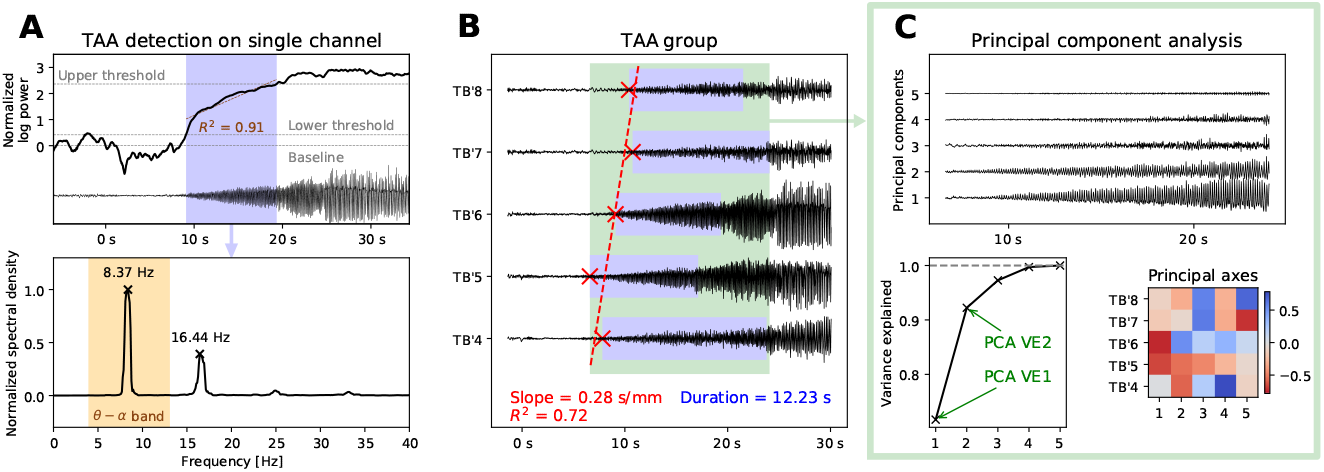
Detection of TAAs and extracting the features of a TAA group. (A) For each channel, we calculate the normalized log-power in the *θ* – *α* band and determine the period of growth (in blue) from the baseline to full seizure activity. We then test if the growth is roughly close to linear (*R*^2^ > 0.75). If so, we check whether the largest peak of the normalized spectral density lies in the *θ* – *α* band and whether all other peaks are its harmonics. (B) When the TAA pattern is detected on four or more consecutive contacts on a single electrode, three features are extracted: the slope and coefficient of determination of the TAA onsets (in red) and average duration of the TAA patterns (in blue). (C) In the TAA interval, the principal component analysis of the SEEG signals is done to extract two more features: the variance explained by the first one and first two components.

From these TAA groups we extracted five features on which our analysis is based: slope of the onset of the TAA patterns and the coefficient of determination as obtained by linear regression, average duration of the TAA patterns, and the variance explained by first two PCA components (Fig. 3B,C). The choice of these features was motivated by the goal of assessing the plausibility of the spreading seizure model. The first three features (slope, *R*^2^, and TAA duration) are informative about the hypothesized seizure spread (or its absence), while the PCA result says more about the internal organization of the seizure activity (such as the fast traveling waves).

Fig. 4 summarizes where and when the TAA patterns were detected, and Tab. 1 summarizes the number of detected TAA groups for the models and the recordings. In the recordings, the TAA pattern was detected on around 2.5% of channels (around 10% of channels where seizure activity was detected), only around one quarter of these were in a TAA group (i.e. more than four TAA patterns on the consecutive contacts of a single electrode). In contrast to that, in all computational models the TAA pattern was detected on majority of contacts with seizure activity (Fig. 4A). That is however to be expected from the models designed to produce the TAA patterns.

**Figure 4:**
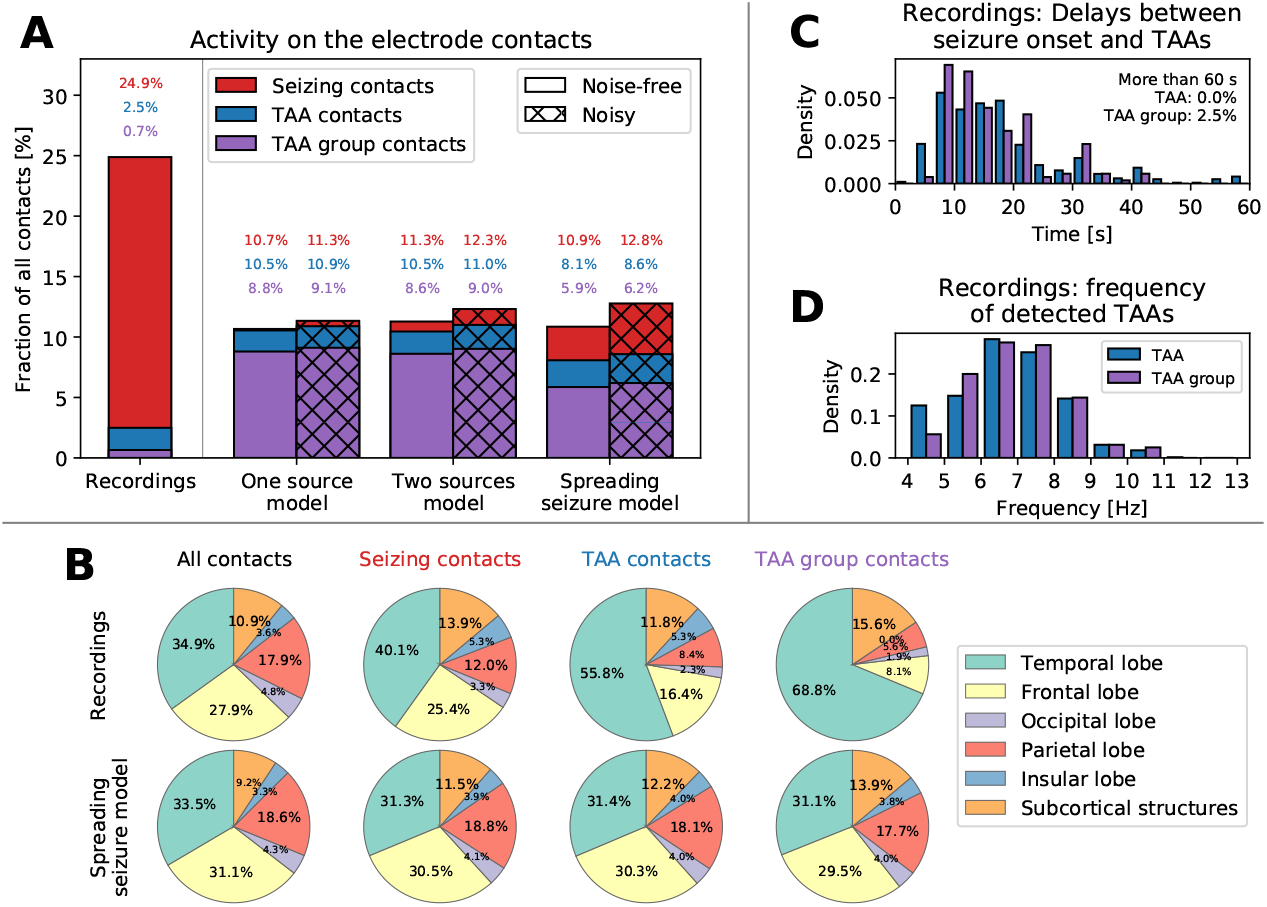
(A) Classification of the contacts based on the recorded/simulated activity. Seizure activity was detected on around 25% of contacts in the recordings and around 10% in the simulations. The TAA pattern was detected on a subset of these seizing contact, and contacts that belong to a TAA group formed even smaller subset. (B) Location of the contacts in the brain. In the spreading seizure model, the TAA patterns are distributed homogeneously in the brain, following the implantation (first column). In the recordings, the TAA patterns occur dominantly in the temporal lobe. (C) Delays of the TAA pattern onset relative to the clinically marked seizure onset in the recordings. Majority of the TAAs appear between eight to twenty second after the seizure onset. (D) Frequencies of the oscillations in the TAA patterns detected in the patient recordings.

**Table 1:**
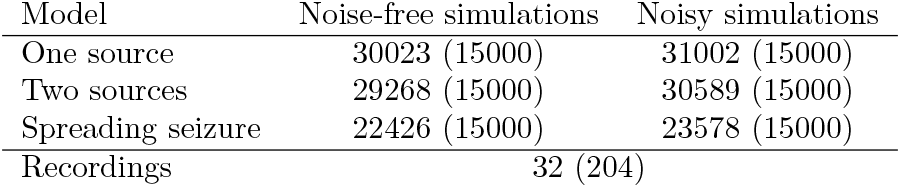
Number of detected TAA groups in the simulated and recorded SEEG. Number in the bracket is the number of simulated or recorded seizures, in which these patterns were searched for.

Analysis of the location of the contacts where the TAA patterns were detected reveal that they dominantly occur in the temporal lobe (Fig. 4B). This observation cannot be explained by a biased electrode implantation - the ratio of the number of contacts where TAA pattern was detected and the number of implanted contacts is higher in the temporal lobe. That is something that the spreading seizure model cannot reproduce. Most of the TAA patterns appear more than eight seconds after the clinically marked seizure onset (Fig. 4C), supporting the hypothesis that TAA pattern is a propagation pattern. The frequency of the oscillations during the TAA patterns is restricted mainly to the interval 5 to 9 Hz (Fig. 4D).

### Features of the recorded and simulated TAA groups

As demonstrated on Fig. 2 (bottom panels), all models can produce activity which at least visually resembles the TAA pattern. To quantify how well the models fit the empirical data, we detected the TAA patterns in the simulated SEEG in the same way as in the patient recordings, and, for each of the detected TAA groups we extracted the five group features (Fig. 3B,C). This procedure gave us for each detected TAA group (in recordings as well as all models) a five-dimensional feature vector, or TAA group “fingerprint”.

Fig. 5A shows the densities of the five group features as observed in the recordings and in the simulations. We highlight several points of interest:

- Short TAAs are more numerous (panel Duration). The empirical distribution of the TAA duration is skewed towards shorter TAAs with duration of 3-7 seconds, and this aspect is reproduced by the spreading seizure model. In contrast, the duration of TAAs in the one and two source model is determined by the prescribed uniform distribution.
- Sequential recruitment is rare even under spreading seizure model (panel Slope). The delayed appearance of the TAA pattern on neighboring contacts would be the most predictive sign of a spreading seizure. Yet, as our model demonstrates, it is a relatively rare event due to the spatial constraint imposed upon the alignment of the electrode with the direction of seizure spread.
- Majority of TAA groups are highly correlated (panel PCA VE1). The trend is reproduced by the two source model as well as the spreading seizure model, although they both predict higher number of highly correlated groups, even in their noisy variations.

**Figure 5:**
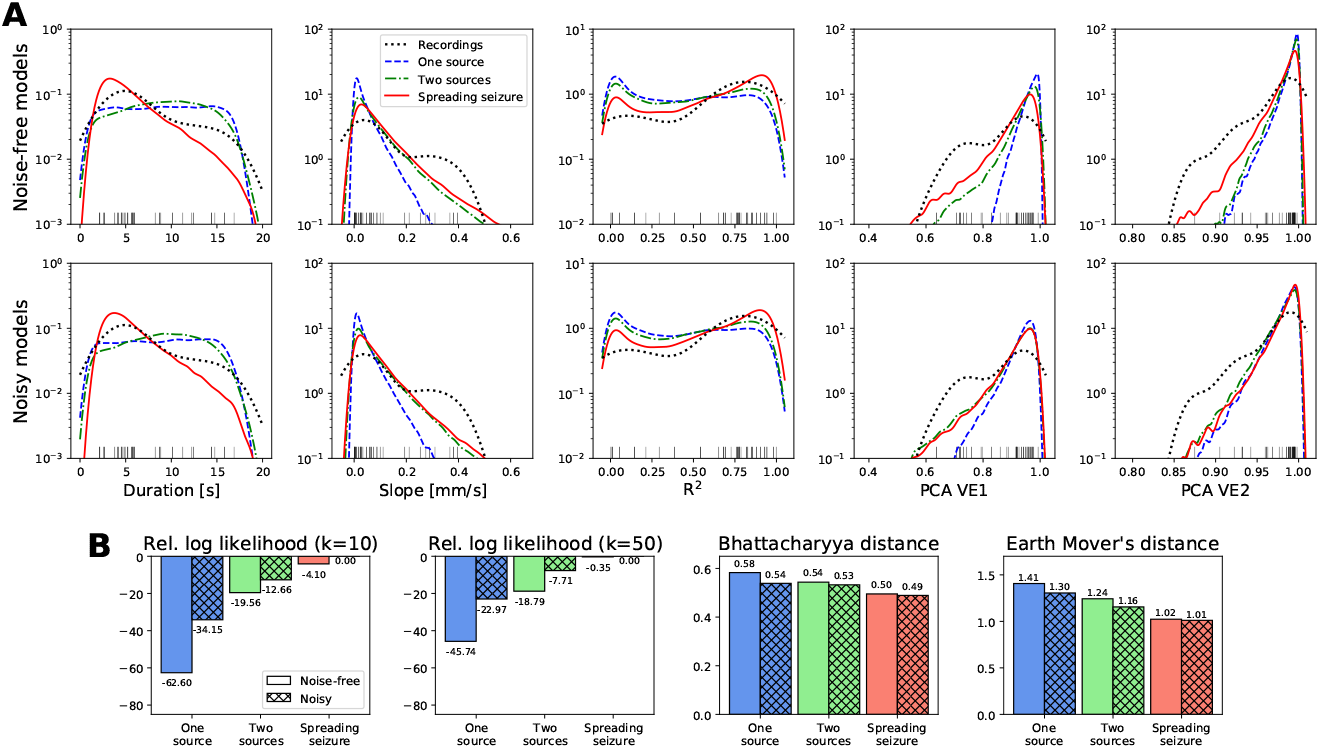
(A) Features of the detected TAA groups in the recordings and simulations. Each panel shows the density of one of the five features of the TAA group (see Fig. 3) obtained from the samples via kernel density estimation for noise-free (top row) and noisy (bottom row) simulations. Each black tick at the bottom edge corresponds to a TAA group in the recordings; these ticks match the density marked by the black dotted line, and are the same in both rows. (B) Quantification of the fit of the models to the empirical data. Each panel shows a measure of a distance of two distributions (empirical and model-generated) in *d*-dimensional space, where *d* = 5 is the number of features. The values of *k*-nearest-neighbor estimation of log-likelihood (more is better) and Bhattacharyya and Earth Mover’s distances (less is better) are shown. For the log-likelihood only the differences (and not the absolute values) are informative; we normalize the values by subtracting the highest value in each chart. To calculate the Bhattacharyya and Earth Mover’s distances, the samples were binned into 1024 uniform bins (each dimension divided into four bins) with limits indicated by the outermost ticks on x-axes in panel A.

### Empirical data are most consistent with the spreading seizure model

Given the “fingerprints” of the TAA groups (i.e. the samples from the distributions on 5-dimensional space of TAA group features) obtained from the recordings on one hand and from the simulations on the other hand, we want to quantify the distance between the recording distribution and all model distributions. Since any single measure of goodness-of-fit has its own advantages and disadvantages, we employ three different measures to assure that the results are robust: *k*-nearest-neighbor approximation of the log-likelihood (with *k* = 10 as well as *k* = 50), Bhattacharyya distance and Earth Mover’s distance (Fig. 5B). The log-likelihood measures how probable are the empirical samples given the model. The Bhattacharyya distance measures the overlap of two probability distributions. For two non-overlapping distributions it gives infinite value, unlike the Earth Mover’s distance, which takes into account how far apart the probability masses are. All measures indicate that the spreading seizure model in its noisy variant fits the empirical distribution of TAA features best among the evaluated models.

### Analysis of the spreading seizure model

Finally, we investigated the behavior of the spreading seizure model in more detail. By performing linear regression between the model parameters and the features of the detected TAAs we demonstrate which parameters affect which features (Fig. 6A and Supplementary Fig. A.8). While all parameters have some effect on most of the features (with the exception of the robust coefficient of determination *R*^2^), some relations are stronger and worth discussing. Unsurprisingly, the strongest relation is between the spread velocity and the TAA duration and the slope of the TAA spread (Fig. 6B): faster spread results in shorter TAA duration and smaller slope of TAA spread.

**Figure 6:**
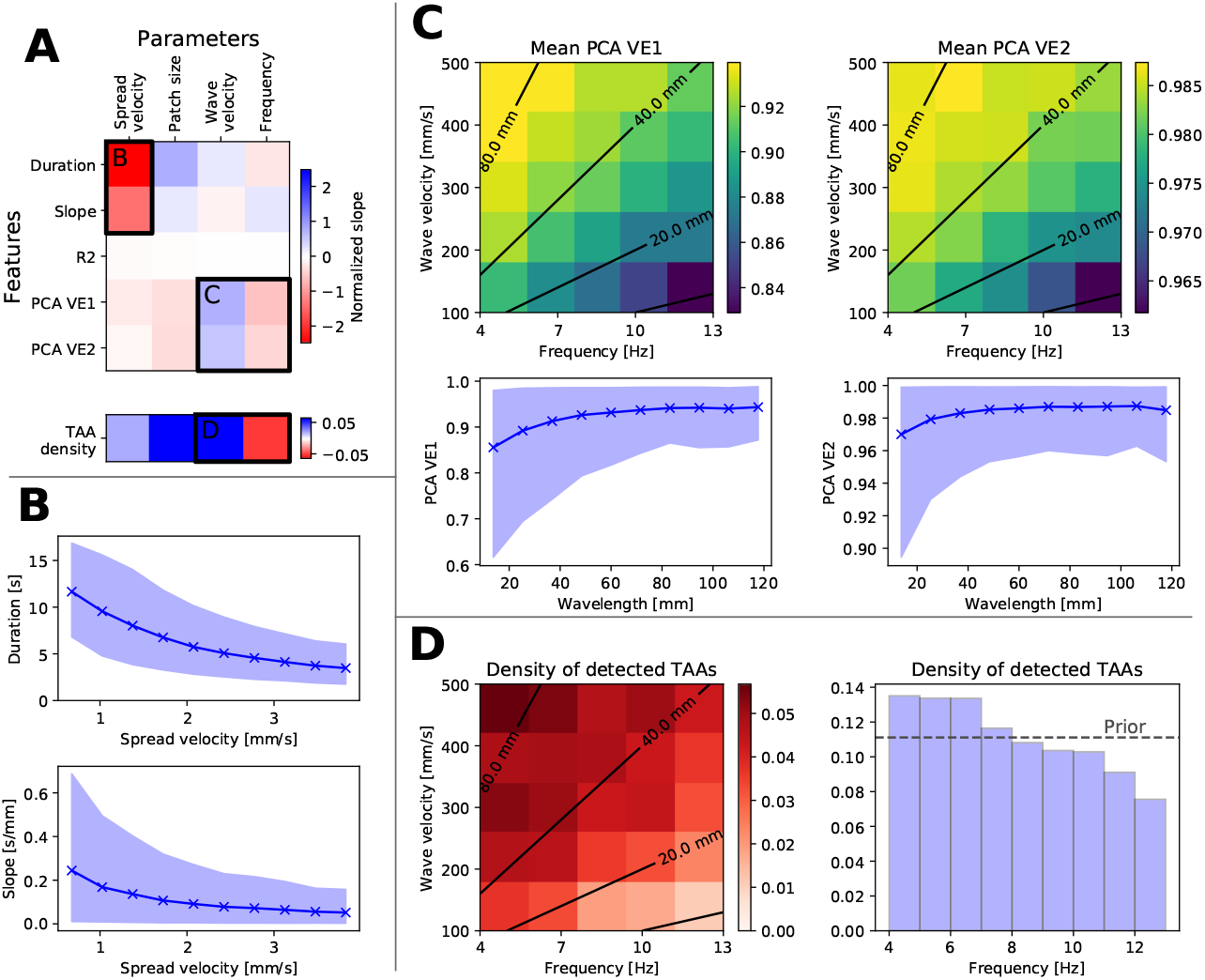
Effects of the model parameters on the features of detected TAAs in the spreading seizure model in the noisy variant. (A) Upper plot shows the slope of linear regression *s_p,f_* between all parameters and features, normalized by the parameter range width *w_p_* and standard deviation *σ_f_* of the feature (*s_p,f_w_p_*/*σ_f_*). Lower plot shows the normalized change in parameter mean for detected TAAs and the prior distribution, 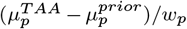, indicating shift of the parameter density to higher/lower values. More saturated blue (red) elements indicate stronger positive (negative) relation. Labeled regions are analyzed in other panels. (B) Relation between the spread velocity and duration of the TAAs and slope of the TAA spread. Solid line and points represent the mean of the features, the shaded area is the 10-90 percentile range. (C) Relation between the frequency and wave velocity and variance explained by the first one and first two components. In the upper plots, black contours indicate the wavelength = wave velocity / frequency. The relation between the features and the wavelength is shown in the lower panels. (D) Relation between the frequency and wave velocity and the density of detected TAAs. The number of detected TAAs decreases with shorter wavelength (left panel) as well as with the frequency alone (right panel).

The role of the wave velocity and frequency of the oscillations is best understood when considered together (Fig. 6C). These two parameters define the wavelength of the fast waves, λ = *u*_wave_/*f*. Longer wavelengths result in larger areas of the seizing patch being synchronized, leading to higher correlations of the simulated signals and thus higher PCA VE1 and PCA VE2 values.

### Higher frequency TAAs are less observed in the spreading seizure model

In the spreading seizure model, the frequency of the oscillations in the generated TAA patterns is determined by the oscillations of the underlying source. However, the model shows that the TAAs with the longer wavelengths are detected more often than the ones with shorter wavelengths (Fig. 6D), presumably again because of the larger areas being synchronized. The same relation holds when looking at the frequency alone - higher frequencies are more rare than the lower frequencies among the detected TAA groups.

This observation indicates that there are two possible explanations of the observed upper limit of the frequencies in the detected TAA patterns (Fig. 4D): either it indeed reflects the restricted range of frequencies of the source oscillations, or, as in our model, it is simply a result of the source to sensor projection which highlights the lower frequency TAAs. In that case, the oscillations caused by traveling waves can occur even at higher frequencies, but may go undetected.

## 3. Discussion

### Summary of the results

In this work we examined the hypothesis that the TAA pattern is caused by seizures spreading across the cortex. The hypothesis ties together the observed SEEG pattern with the observations of the spreading seizures (Schevon et al., 2012; Martinet et al., 2015) and fast traveling waves during seizures (Smith et al., 2016; Martinet et al., 2017) in fashion consistent with recent models of seizure dynamics (Proix et al., 2018; Liou et al., 2019). We analyzed the SEEG recordings of epileptic seizures collected from a cohort of fifty patients and compared how the predictions of three different models fit the data. Among the tested models, the predictions of the spreading seizure model in the noisy variation were closest to the observed data. The model with realistic parameters is consistent with most features of the observed TAA patterns, including a) the duration of the TAA patterns of several seconds (Fig. 5A, panel Duration), b) rarely observed sequential recruitment of neighboring contacts (Fig. 5A, panel Slope), or c) the high percentage of TAAs with highly correlated signals (Fig. 5A, panel PCA VE1). In addition, the model may provide an explanation for the upper limit of the TAA frequency range (Fig. 6D). There are however several observations that the spreading seizure model cannot explain, mainly why are the TAA patterns disproportionately often observed in the temporal lobe (Fig. 4B).

### Relation of TAA and spreading seizures

Our results show that the spreading seizure model is the most plausible explanation when looking at all detected TAA patterns as a group. That, however, does not mean that it is the best explanation for any single TAA even among the limited selection of models we have evaluated. One thus cannot conclude that all TAA occurrences are caused by spreading seizures; other explanations might be plausible for any individual case. Indeed, TAA patterns were observed not only in the sites of seizure propagation, but also within the epileptogenic lesions (Lagarde et al., 2016), where the explanation by spreading seizure might be questioned.

One might consider fitting the models for individual TAA patterns using the information about the geometry of the cortex in the vicinity of the electrode of interest in order to obtain the parameter values best explaining the individual pattern. These parameters might include, in case of the spreading seizure model, the velocity and direction of the spread or the location of the seizure patch. The fitted models could then be compared to establish the probable cause of each pattern. This effort would however be complicated by factors shared by many inverse problems, mainly large degree of freedom, ill-posedness and issues linked to identifiability (Grech et al., 2008; Daunizeau et al., 2011). To deal with such issues, strong regularization of the model corresponding to strong assumptions about the nature of the source activity would probably be needed, limiting the usefulness of the results.

### Frequency of occurrence of TAA patterns

In the seizure recordings the TAA pattern was detected in 10% of the channels with seizure activity. That is partly a consequence of the utilized detection mechanism: on purpose we have used relatively strict criteria to define what is a TAA pattern in order to get TAA patterns from which the features can be robustly extracted. If, instead, we opted to classify all channels according to the electrographic patterns into several distinct categories, more questionable patterns could have been classified as TAA patterns as well and the percentage could be higher. Indeed, Schiller et al. (1998) found that in the sites of seizure propagation in mesiotemporal lobe, 53% of early patterns consisted of “rhythmic round theta and delta activity” correlated with activity at the onset site, possibly related to the TAA pattern as we defined it here. In the sites of seizure propagation in neocortex the proportion was even higher (82%). Also Perucca et al. (2014) states that the TAA pattern was found in the propagation sites in more than half of the seizures.

### Thalamic involvement

In this work we do not explore the biological or dynamical mechanisms of the traveling waves, only its manifestations on the intracranial recordings. While the traveling waves might be generated and sustained purely by the cortex as predicted by some models of seizure dynamics (Proix et al., 2018; Liou et al., 2019), they might be also driven by or arise from interactions with other structures. One such structure could be the thalamus. Thalamus was for long implicated in the generation of rhythmic spike-and-wave discharges in absence seizures (Destexhe, 2008; Avoli, 2012) but evidence exists for its involvement in focal seizures as well (Guye et al., 2006; Blumenfeld et al., 2009; He et al., 2017). Interestingly, experimental and computational evidence show that thalamus can support propagating waves, both in isolation or as a part of a coupled thalamocortical system (Muller and Destexhe, 2012). Its involvement in temporal lobe epilepsies (Guye et al., 2006) could explain the predominant occurrence of the detected TAA patterns in the temporal lobe.

### Modeling of TAA patterns in large scale brain models

Although the forward problem, aka map between source and sensor space, is clearly acknowledged as a contributor to the difficulty in performing correctly model inversion, often such recognition remains academic. Clinically, SEEG electrode contacts are regularly identified synonymously with brain region locations (electrode contacts A1-A2 for right hippocampus, B1-B2 for right amygdala, etc.), although there is a well-established variability in terms of surgical implantation and subsequent clinical interpretation, with consequences for surgery success rate and scientific reproducibility, in case the data are used for research. Our contribution reaches beyond this known spatial confound of the forward solution leading to misidentification of the epileptogenic zone due to displacement; in fact, we render the forward problem spatiotemporal by demonstrating how well-established clinical marker, the TAA pattern, can be mimicked through an appropriate network dynamics. This demonstration highlights the importance of the necessity to integrate quantitative neuroinformatics tools such as The Virtual Brain (Sanz Leon et al., 2013) in the workflows of interpretation and decision making in epilepsy research.

Epilepsy research is particularly deeply connected to nonlinear dynamics. One of the few fundamental discriminants of nonlinear dynamics is sudden qualitative changes in behavior, which are technically described by bifurcations (Haken, 1977; Kuznetsov, 1998; Izhikevich, 2010). They are powerful tools of modeling, because the behavior of any dynamic system can be smoothly mapped upon a set of canonical equations (so-called normal form) that is representative for the transition. In epilepsy, there is large evidence that the majority of seizure onset and offset transitions can be understood as bifurcations (Touboul et al., 2011; Jirsa et al., 2014; Bernard et al., 2014), although also other forms of transitions have been recognized, for instance in absence seizures so-called false bifurcations, in which significant changes occur rather smoothly than discretely (Marten et al., 2009). Bifurcation analysis permits the construction of a seizure taxonomy based on purely dynamic features (Jirsa et al., 2014; Saggio et al., 2017), which guides further research and may, for instance, lead to the discovery of novel attractors (Houssaini et al., 2015). From the perspective of bifurcations, the TAA patterns would be classified as a dynamical system undergoing a supercritical Hopf bifurcation, whose distinguishing feature is the gradually increasing amplitude of the oscillations after crossing the bifurcation (Jirsa et al., 2014; Saggio et al., 2017). As we have demonstrated here, the TAA pattern can also be produced by spreading seizures with fast traveling waves. It was shown that these features can be generated using the spatially extended Epileptor model (Proix et al., 2018), which uses the saddle-node bifurcation to initiate the seizure. This misidentification has two consequences: first, the extent of the oscillation source in a spreading seizure model changes over time, thus a simple spatial inversion assuming a static source is not appropriate and could lead to a misidentification of the epileptogenic zone; second, the bifurcation type may be incorrectly inferred, which has important consequences for the behavior of the dynamical system, such as the response to external stimulation, responses to local interventions via drug administrations and difference in network propagation. For instance, stimulation of a dynamic system close to a supercritical Hopf bifurcation will not be able to trigger a seizure, whereas stimulation close to a saddle-node bifurcation generally will. Such is a consequence of the different properties of multistability following from each bifurcation. The correct identification of the bifurcation type is thus critical and clinically consequential, because the different system properties are linked to therapeutic interventions. As the discovery of novel forms of network modulation (other than resective surgery) is currently an active field of research in epilepsy, the integration of spatial and spatiotemporal consequences of realistic forward modeling is an important element to be integrated in the future workflows.

### Limitations of the study

The results have to be interpreted within the limitations of the study. Importantly, it is the sample size. Even with a large cohort of fifty patients and multiple seizure recordings for each patient, only 32 TAA groups were detected in the recordings, and the model comparison was based on these 32 samples. Although it is only the model comparison that is affected by the small sample size and the model predictions are independent, the conclusions need to be considered in light thereof.

Further limitation is that the results of the model comparison inevitably depend on exact form of the models and the choice of the ranges of the model parameters. We have not attempted to fit the parameters to the observed data, instead, we chose the parameters distributions based on the experimental observations in the literature, and our choices could be questioned. We therefore advise not too interpret the results in an overly formal way such as evaluating Bayes factors leading to the statements on the strength of evidence in favor of one or the other model (Kass and Raftery, 1995). The main goal of the work was not to select the best model from the candidate models; such task is not even necessary. Rather it was to gain insight about the behavior, predictions, and shortcomings of the spreading seizure model when compared to the empirical data. The other models served us mainly as a baseline in this endeavor.

## 4. Methods

### 4.1 Patient data

In this study we have used imaging and electrographic data from a cohort of 50 patients who underwent a clinical presurgical evaluation (Tab. A.3). The clinical evaluation was described in detail before (Proix et al., 2017). The T1-weighted images (MPRAGE sequence, repetition time = 1900 ms, echo time = 2.19 ms, 1.0 × 1.0 × 1.0 mm, 208 slices) were obtained on a Siemens Magnetom Verio 3T MR-scanner. The patients were implanted with multiple stereotactic EEG electrodes. Each electrode has up to 18 contacts (2mm long a 0.8mm in diameter), which are either uniformly placed on the electrode, separated by 1.5 mm from each other, or placed in groups of five, which are separated by 9 mm. The SEEG was recorded by a 128 channel Deltamed™ system using at least 256 Hz sampling rate. The recordings were band-pass filtered between 0.16 and 97 Hz by a hardware filter. After the electrode implantation, a CT scan of the patient’s brain was acquired to obtain the location of the implanted electrodes. The patient signed an informed consent form according to the rules of local ethics committee (Comité de Protection des Personnes Sud-Méditerranée I).

### 4.2 Cortical surface reconstruction

The brain anatomy was reconstructed from the T1-weighted images by FreeSurfer v6.0.0 (Fischl, 2012) using the *recon-all* procedure. Afterwards, the triangulated surface used in this study was obtained by taking the midsurface of the pial surface and white matter-gray matter interface, i.e. the surface lying halfway between these surfaces. The position of the contacts in the CT scan with implanted electrodes was marked using the GARDEL software (Villalon et al., 2018), and then transformed to the T1 space using the linear transformation obtained using the linear registration tool FLIRT from the FSL toolbox (Jenkinson et al., 2002).

### 4.3 Computational models

The models define the source activity on the subject’s cortex Ω, and its projection on the sensors implanted in the brain. Specifically, the models prescribe the seizure activity on the recruited part Ω_*r*_(*t*) of the excitable patch Ω_*e*_ of the cortex, Ω_*r*_(*t*) ⊆ Ω_*e*_ ⊆ Ω. On the rest of the cortex not recruited in the seizure activity only noise is prescribed.

#### One source model

In the one homogeneous source model, the whole excitable patch is recruited at time *t*^0^,

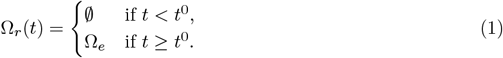

The excitable patch Ω_*e*_ is in each simulation selected randomly by choosing a center point on the cortex Ω which is placed closer than 15 mm from any of the electrode contacts. Then the patch size is chosen from its prescribed range (Tab. 2), and the patch is extended from the initial point until the desired size is reached. The source activity at point ***x*** on the cortical surface and time *t* is given by

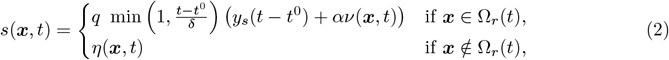

where

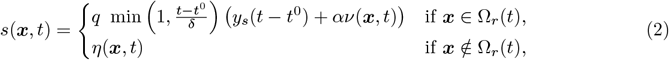

is a triangle wave with frequency *f*, with power normalization constant 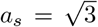. Next, *η*(***x***, *t*) and *ν*(***x***, *t*) is the background and seizure noise, described below, with the latter present only for the noisy simulations, i.e. *α* = 0 for noise-free and *α* = 1 for noisy simulations. Finally, *δ* is the duration of the onset pattern and *q* is a scaling coefficient (Tab. 2).

**Table 2:**
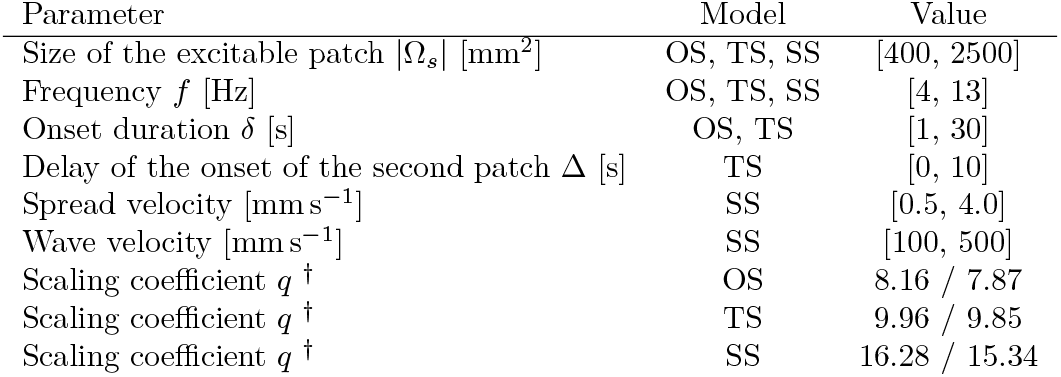
Ranges and values of model parameters. Model abbreviations: OS - One Source, TS - Two Sources, SS - Spreading seizure. ^†^ For noise-free / noisy simulations.

| Parameter | Model | Value |
| --- | --- | --- |
| Size of the excitable patch $\vert \Omega_s\vert $ [mm<sup>2</sup>] | OS, TS, SS | [400, 2500] |
| Frequency $f$ [Hz] | OS, TS, SS | [4, 13] |
| Onset duration $\delta$ [s] | OS, TS | [1, 30] |
| Delay of the onset of the second patch $\Delta$ [s] | TS | [0, 10] |
| Spread velocity [mm s<sup>-1</sup>] | SS | [0.5, 4.0] |
| Wave velocity [mm s<sup>-1</sup>] | SS | [100, 500] |
| Scaling coefficient $q$ <sup>†</sup> | OS | 8.16 / 7.87 |
| Scaling coefficient $q$ <sup>†</sup> | TS | 9.96 / 9.85 |
| Scaling coefficient $q$ <sup>†</sup> | SS | 16.28 / 15.34 |

#### Two sources model

In the two homogeneous sources model, there are two excitable patches which are recruited with delay Δ, so that 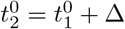 and

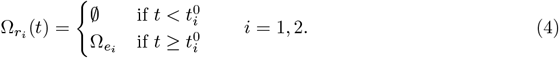

The excitable patches Ω_*e*_1__ and Ω_*e*_2__ are selected in a similar way as in the one source model. First, the patch centers are randomly chosen so that they lie closer than 15 mm from any of the electrode contacts and also closer to each other than 10 mm. Then the patch size is chosen, and each of the patches is expanded to half of its value. The source activity is given by

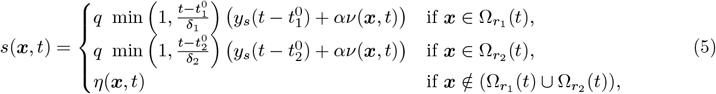

with the waveform *y_s_* given by Eq. 3, background and seizure noise *η*(***x***, *t*) and *ν*(***x***, *t*) described below, and onset duration *δ*_1_ and *δ_2_* and scaling coefficient *q* as in Tab. 2.

#### Spreading seizure model

With the propagating seizure model, the excitable patch is recruited sequentially,

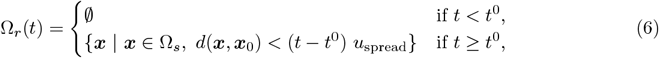

where *d*(***x, x***_0_) is the geodesic distance of the point ***x*** from the seizure origin ***x***_0_ and *u*_spread_ is the velocity of seizure spread. The excitable patch is created the same way as in the one source model, and the seizure origin ***x***_0_ is selected randomly from all points on the excitable patch. The cortical activity is prescribed as

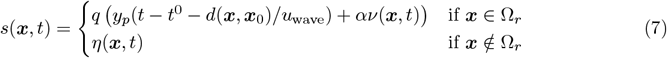

where

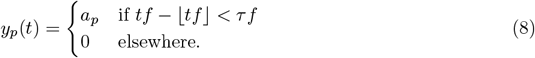

is a pulse wave with duty cycle *τ* = 0.25 and frequency *f*, 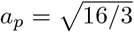 is the power normalization coefficient, the parameter *u*_wave_ is the velocity of the fast waves traveling across the recruited part of the excitable patch, *q* is the scaling coefficient, and *η*(***x***, *t*) and *ν*(***x***, *t*) is the background and seizure noise.

#### Noise

The noise in the simulations consist of the background noise *η*(***x***, *t*) present in the regions not recruited into seizure activity, and (in case of noisy simulations) the seizure noise *ν*(***x***, *t*). Both the background and seizure noise are modeled as pink (i.e. 1/*f*) noise with power equal to one. The seizure noise is modeled as spatially correlated with kernel *k*(***x, x′***) = exp (−*d*(***x, x′***)/*l*) with *l* = 10 mm, *d*(***x, x′***) is the geodesic distance of two points on the cortical surface. For reasons of computational efficiency, the background noise of the whole cortex is not modeled as spatially correlated, instead, the cortex is divided into patches of average area 100 mm^2^, and the same noise time series is used for all points in one patch. The division is performed by randomly selecting appropriate number of seed vertices and then expanding all patches until the whole cortex is covered.

The scaling coefficient *q* was set experimentally so that the normalized log power of the simulated SEEG would have comparable maximal values as the normalized log power of the recorded signals. Specifically, for each model and for both noisy and noise-free simulations, sixty simulations with randomly chosen parameters were performed with fixed 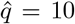. Then the 80th percentile of log power normalized to preictal levels was calculated for all channels (across time), and 95 percentile *p*_sim_ was again calculated across all channels and simulated seizures. The same value *p*_rec_ was calculated for the recorded seizures, and the scaling coefficient was updated, 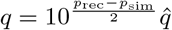.

#### SEEG projection

In the human cortex, the most numerous neuron type is the pyramidal cell. Due to their geometrical structure with long dendrites oriented perpendicularly to the cortical surface, they might be well represented as electrical dipoles (Buzsáki et al., 2012). Following this idea, we assume that each point on the cortical surface acts as electrical dipole. In the spatially continuous formulation the local field potential measured by the electrode contact at point ***x***_*s*_ generated by the source activity *s*(***x***, *t*) on the surface Ω is given by

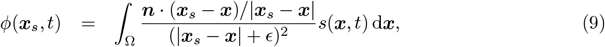

where ***n*** is the outward oriented normal of the surface, and the constant *ϵ* = 1 mm is added to prevent the singularities at the surface. In the discretized version using the calculated solution on a triangulated surface the formula is instead

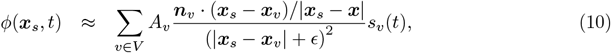

where *V* is the set of all vertices on the triangulated surface, *A_v_* is the area associated with a vertex (calculated as one third of the sum of areas of neighboring triangles), ***n***_*v*_ is the outwards oriented normal of a vertex (calculated as a weighted average of the normals of neighboring triangles), ***x***_*v*_ is the position of the vertex, and *s_v_*(*t*) is the activity at the vertex prescribed by one of the above models (Eq. (2), (5), or (7)).

### 4.4. TAA pattern detection

The occurrences of the TAA pattern in all channels of the recorded (or simulated) electrographic signals are detected in two steps. In the first step, we look for the temporal interval where the power in the *θ* – *α* band grows from the preictal baseline to full seizure activity. In the second step, we check that this tentative interval satisfies further conditions on the TAA pattern, namely that the oscillatory activity is dominated by single frequency oscillations in the *θ* – *α* range and the growth is close to linear.

The first step is implemented as follows. The power in the *θ* – *α* band (4 to 13 Hz) of the SEEG signals in monopolar representation is calculated using the multitaper method (time bandwidth = 2.0, number of cycles = 8). The *θ* – *α* log-power *LP_θ–α_* is then normalized to preictal baseline (calculated from sixty seconds preceding the clinically marked seizure onset) so that 〈*LP_θ–α_*〉_preictal_ = 0. The channel is marked as *seizing* if the 90-th percentile of the power *P*_90_ is at least 30-times larger than the baseline. If it is not seizing, than it is also marked as not TAA. Otherwise, the limits of the tentative interval of the TAA pattern [*t^o^, t^t^*] are set as *t^t^* = min{*t* | *LP_θ–α_*(*t*) < *k*_2_} and *t^o^* = max{*t* | *t* < *t^t^*, *LP_θ–α_*(*t*) > *k*_1_}, where the lower threshold is set to *k*_1_ = 0.15 *P*_90_ and upper threshold to *k*_2_ = 0.85 *P*_90_. This procedure implies that this tentative TAA interval can be preceded by any activity as long as its *θ* – *α* log-power does not cross the *k*_2_ threshold; the TAA patterns that we detect can thus appear after abnormal electrographic activity of low power or outside of the *θ* – *α* band.

Next, we check that the signal in the tentative interval satisfies our criteria on TAA patterns. First, we apply the linear regression on the log-power over time, and check that the coefficient of determination is sufficiently high, *R*^2^ > 0.75. Second, we calculate the power spectral density of the signal in the tentatively determined interval in the range 1 to 100 Hz, we flatten the spectrum by multiplying it by the frequencies, and we normalize it so that the maximum is equal to one. We then detect the peaks in the spectrum (minimum peak height 0.25, peak distance 2 Hz). The pattern is confirmed to be the TAA pattern if the frequency of the largest peak *f*_0_ lies in the *θ* – *α* range (i.e. 4 to 13 Hz), and if all other peaks *f_i_* are harmonics of *f*_0_ (with tolerance 0.15 *f*_0_, i.e |*f_i_* – *kf*_0_| < 0.15 *f*_0_ for some *k* = 1, 2,…).

When the TAA pattern is detected on four or more neighboring contacts of a single electrode, five features are computed for the group. Two are obtained from linear regression of the TAA onset times *t^o^* w.r.t. the position on the electrode: *slope* and the coefficient of determination *R*^2^. Next is the average *duration* of the TAA pattern, i.e. 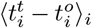, where *i* indexes the contacts in the group. The last two features are determined by performing a principal component analysis of the recorded (or simulated) signals in the interval 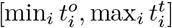, and calculating the variance explained by the first one (*PCA VE1*) and first two (*PCA VE2*) components.

### 4.5 Measures of goodness-of-fit

We use three measures to quantify the fit between the features of the TAA patterns detected in the recordings and in the simulated data sets. In the following paragraphs, we assume that from the recordings we have *n d*-dimensional samples 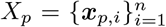 from the probability distribution *P* with unknown probability density function *p*, and that we have *m* model-generated samples 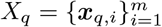 from the distribution *Q* with probability density function *q*.

#### Log-likelihood

The log-likelihood measures how likely are the samples *X_p_* under the hypothesis represented by the probability distribution *Q*,

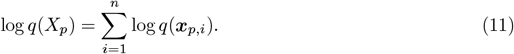

The probability density function *q* is however not known, and we approximate it with its *k*-nearest-neighbor estimation,

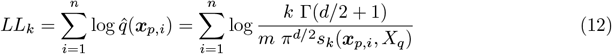

where Γ is the gamma function and *s_k_* is the distance from *x_p,i_* to its *k*-nearest-neighbor among *X_q_*.

#### Bhattacharyya distance

The Bhattacharyya distance (Kailath, 1967) measures the overlap of two probability distributions. It relies on the binning of the samples. Given the probabilities *p_j_* and *q_j_* in *j* = 1…*n_b_* bins, the distance is defined as

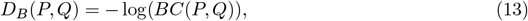

where 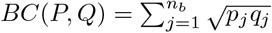 is the Bhattacharyya coefficient.

#### Earth mover’s distance

Intuitively, Earth mover’s distance (Rubner et al., 2000) corresponds to minimal amount of work needed to transport the mass from one distribution to another. Again, we need to bin the samples into bins to obtain the probabilities *p_j_* and *q_j_* in *j* = 1… *n_b_* bins. Then the Earth Mover’s distance is defined as

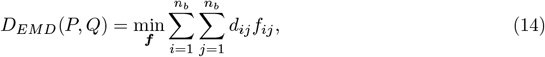

where *d_ij_* is the Euclidean distance between the centers of *i*-th and *j*-th bin. The distances along each dimensions are normalized by the standard deviation of the feature values in the recording samples. The cost function is minimized over all possible flows 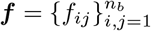, subject to following constraints:

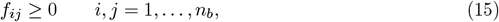

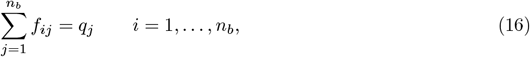

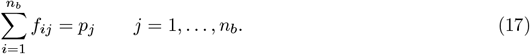

The constraint (15) guarantees that only positive amount of mass is transported, while (16) and (17) limit the amount of mass transported from and to a single bin to the amount given by the distributions. The optimization is implemented using the OR-Tools package (Perron and Furnon, 2019).

## Author contributions

VS and VJ designed the study. MG, FB, and JS acquired and collected the data. VS performed the study. VS, JS, MG, FB, and VJ wrote the manuscript.

## Acknowledgements

The authors wish to acknowledge the financial support of the following agencies: the French National Research Agency (ANR) as part of the second “Investissements d’Avenir” program (ANR-17-RHUS-0004, EPINOV), European Union’s Horizon 2020 Framework Programme for Research and Innovation under the Specific Grant Agreement No. 785907, PHRC-I 2013 EPISODIUM (grant number 2014-27), the Fondation pour la Recherche Médicale (DIC20161236442) and the SATT Sud-Est (827-SA-16-UAM) for providing funding for this research project.

## Data availability

The patient data sets cannot be made publicly available due to the data protection concerns.

## Code availability

The code will be made available upon publication.

## Appendix A. Supplementary material

**Figure A.7:**
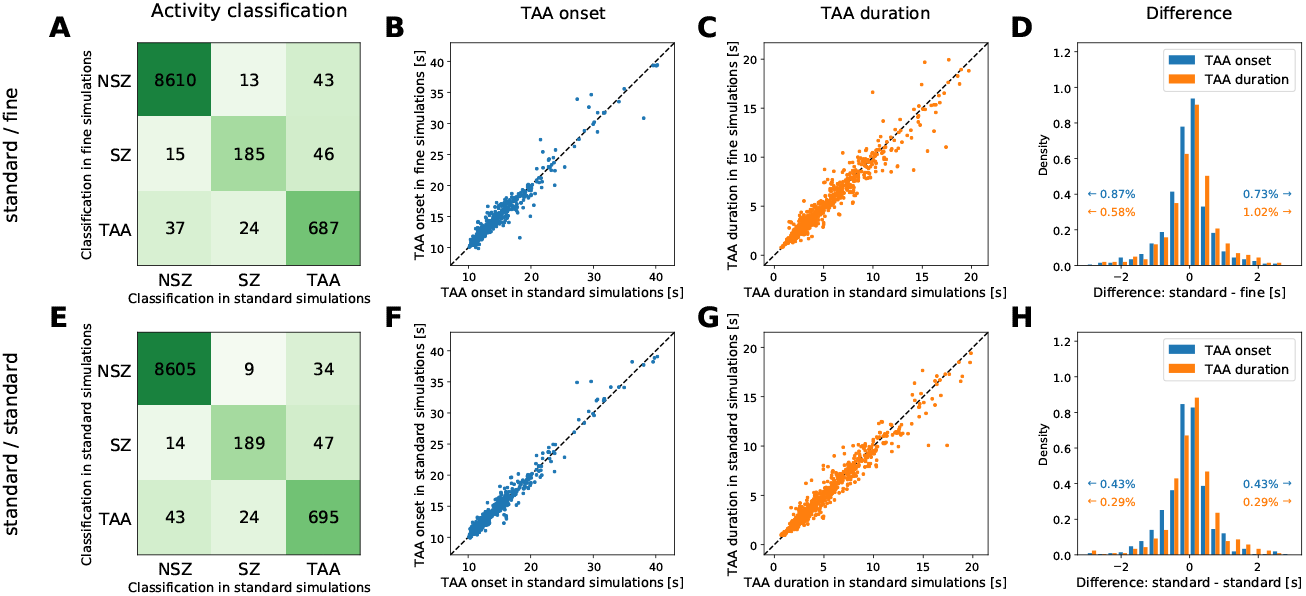
Study of the mesh dependence. (A-D) Sixty simulations of spreading seizure in the noise-free variant of the model were performed on the standard triangulation (described in the main text) and on refined triangulation of the cortex, obtained by splitting every existing triangle into four. The simulation parameters were chosen randomly as in the main text, but were kept between the simulations on standard and fine triangulations, so that the results are directly comparable. The background noise was however different between the simulations on standard and fine triangulation. (A) The activity of the simulated SEEG signals were classified into non-seizing (NSZ), seizing but not TAA (SZ), and TAA (see Methods). (B) For the signals classified as TAA in both standard and fine simulations, the determined time of TAA onset is compared. Perfect fit would lie on the diagonal marked by black dashed line. (C) Same as B but for the determined duration of TAA pattern. (D) Histogram of the differences of the TAA onset times and durations from panels B and C. The range is clipped for visualization, amount of clipped values is shown in the inset text. (E-F) To assess the influence of the background noise, second set of simulations on the standard triangulation was performed. The parameters of the simulations were again kept the same as in the first set, only with the background noise changed. Panels are equivalent to panels A-D, showing the fit between the two sets of simulations on standard triangulations. Comparison between the first and second row of the figure indicates that the differences between the simulations on standard and fine triangulations are caused mainly by the stochastic background noise, since they are present also for the simulations on the same triangulations. The level of mesh refinement thus does not introduce differences of higher order of magnitude.

**Figure A.8:**
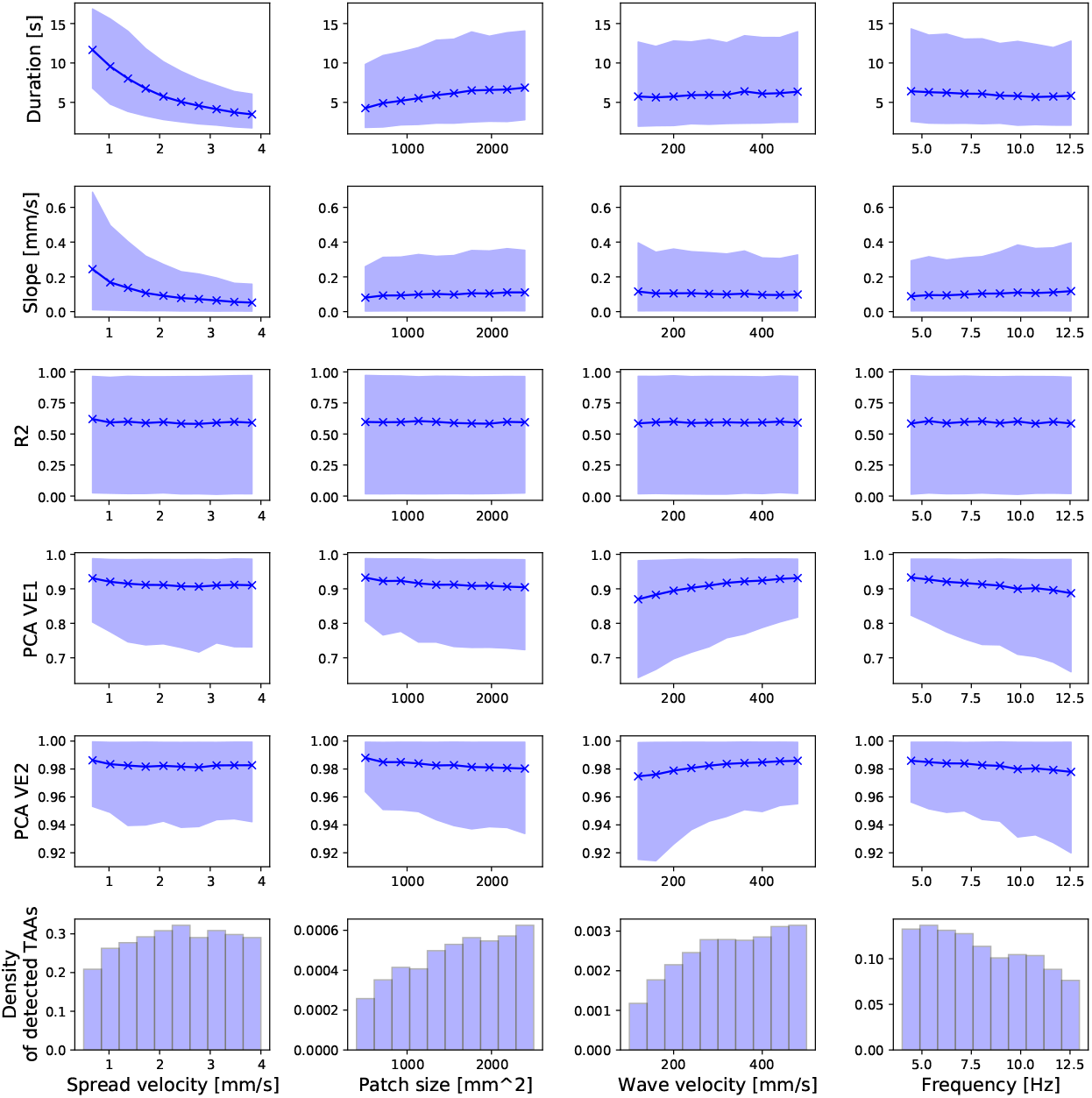
Effects of the parameters in the spreading seizure model in the noisy variant, visualizing the relations on Fig. 6A. Each panel shows the relation between one parameter and one feature. Solid line and points represent the mean of the features, the shaded area is the 10-90 percentile range. The last row shows the histogram of the parameters among the detected TAAs.

**Table A.3:**
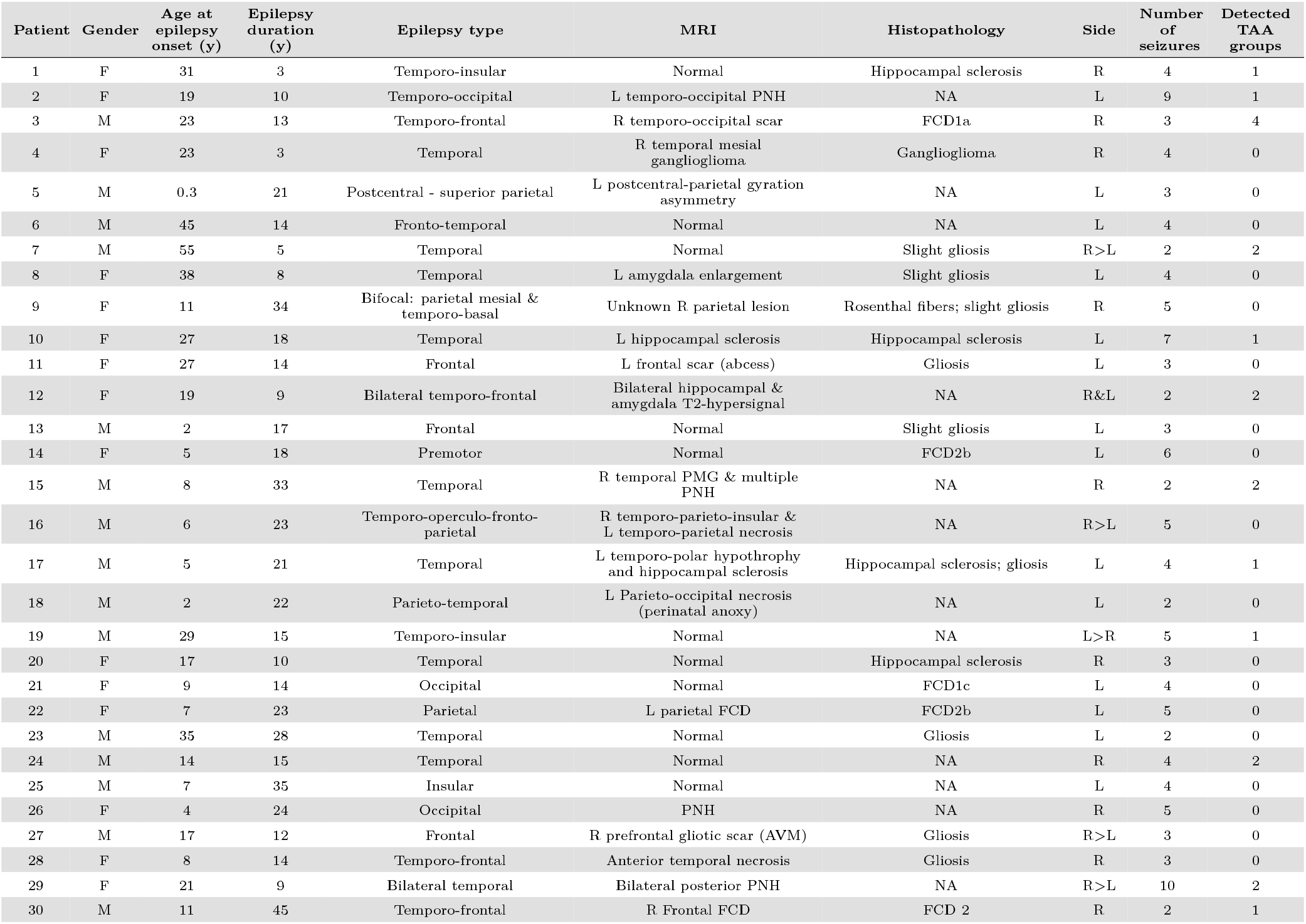
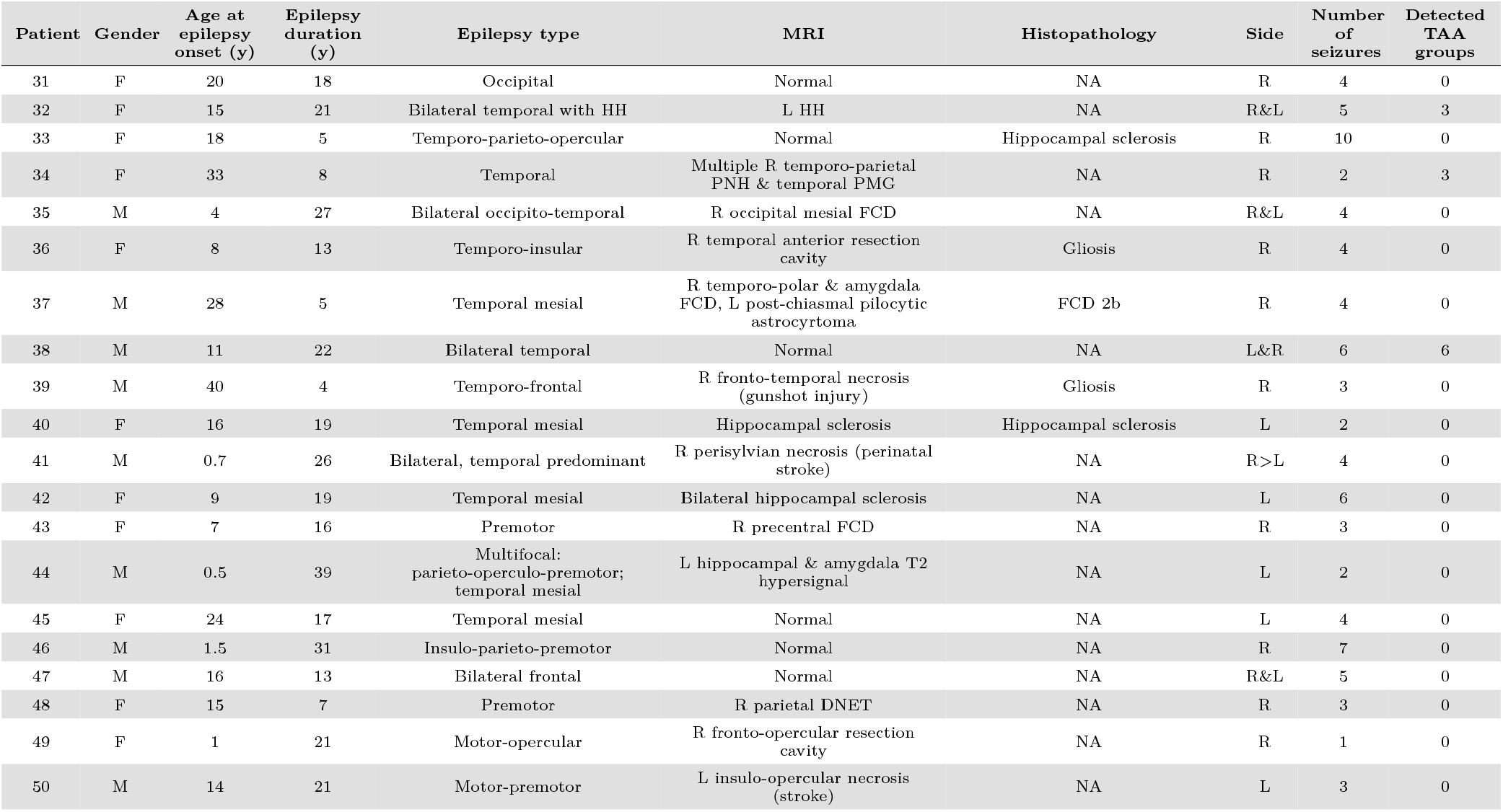
Patient table. Abbreviations: AVM, arteriovenous malformation; DNET, dysembryoplastic neuroepithelial tumor; FCD, focal cortical dyplasia; HH, hypothalamic hamartoma; L, left; NA, not applicable; PMG, polymicrogyria; PNH, periventricular nodular heterotopia; R, right.

